# Rapid Biomarker Screening of Alzheimer’s Disease by Interpretable Machine Learning and Graphene-Assisted Raman Spectroscopy

**DOI:** 10.1101/2021.06.03.446929

**Authors:** Ziyang Wang, Jiarong Ye, Kunyan Zhang, Li Ding, Tomotaroh Granzier-Nakajima, Jeewan Ranasinghe, Yuan Xue, Shubhang Sharma, Isabelle Biase, Mauricio Terrones, Se Hoon Choi, Chongzhao Ran, Rudolph E. Tanzi, Sharon X. Huang, Can Zhang, Shengxi Huang

## Abstract

As the most common cause of dementia, the study of Alzheimer’s disease (AD) faces challenges in terms of understanding the cause, monitoring the pathogenesis, and developing early diagnosis and effective treatment. Rapid and accurate identification of AD biomarkers in the brain is critical to provide key insights into AD and facilitate the development of early diagnosis methods. In this work, we developed a platform that enables a rapid screening of AD biomarkers by employing graphene-assisted Raman spectroscopy and machine learning interpretation in AD transgenic animal brains. Specifically, we collected Raman spectra on slices of mouse brains with and without AD and used machine learning to classify AD and non-AD spectra. By contacting monolayer graphene with the brain slices, the accuracy was significantly increased from 77% to 98% in machine learning classification. Further, using linear supporting vector machine (SVM), we identified a spectral feature importance map that reveals the importance of each Raman wavenumber in classifying AD and non-AD spectra. Based on this spectral feature importance map, we identified AD biomarkers including Aβ and tau proteins, and other potential biomarkers, such as triolein, phosphatidylcholine, and actin, which have been confirmed by other biochemical studies. Our Raman-machine learning integrated method with interpretability is promising to greatly accelerate the study of AD and can be extended to other tissues, biofluids, and for various other diseases.

## INTRODUCTION

Alzheimer’s disease (AD), a progressive disorder of the brain that causes memory losses and damages other brain functions, is the most common cause of dementia.^1^ By 2020, about 44 million people worldwide have been diagnosed with AD.^2^ Despite the prevalence, the causes of AD are still not fully understood. Studying biomarkers related to AD greatly accelerates the understanding of the disease and can lead to new treatment against dementia.^3, 4^ Three biomarkers, T-tau, P-tau, and Aβ_42_, have been identified and confirmed in the cerebrospinal fluid that are strongly associated with AD and could be used as progression markers in developing drugs.^5^ To detect the AD-associated biomarkers in the brain, various imaging techniques have been developed, such as magnetic resonance imaging (MRI) and positron emission tomography (PET).^6–8^ However, MRI and PET are costly and time-consuming while they still lack specific molecular information.^6, 7^ Other biosensing methods such as surface plasmon resonance biosensors and field-effect transistors offer specific information on the optical or electronic properties of the analyte ^8–10^ thus are insufficient to gain comprehensive insights into the biomarkers of AD. Recently, spectroscopy-based detection of AD biomarkers via immunoassay and fluorescence on blood and cerebrospinal fluid (CSF), has been intensively investigated in preclinical stages,^11–14^ but they are not label-free which prevent the discovery of novel biomarkers. New methods to rapidly screen and identify potential AD biomarkers from a huge number of candidate molecules are still urgently needed.

Raman spectroscopy is a non-destructive and label-free molecular sensing method. By exciting the samples with a monochromatic laser and collecting the inelastically scattered signal from the analyte, the obtained Raman spectrum provides the fingerprints of the analyte. Additionally, it offers high multiplexity and high specificity due to the multiple and extremely narrow Raman peaks. Since Raman spectroscopy provides a desirable approach with rapid diagnosis, it has been utilized to investigate AD in terms of diagnosing AD with Lewy Bodies in blood plasma,^15^ classifying early pathological states of AD with brain hippocampus regions,^6^ imaging amyloid plaques in brain tissues,^16^ *etc.* Despite the specificity, multiplexity, and rapid diagnosis, the interpretation of the Raman signals in complex bio-samples is challenging. Although spectral comparison and principal component analysis (PCA) have been employed in Raman spectral analysis, molecule identification is unreliable when the intra-class spectral variation is too high.^17–21^

In recent years, machine learning has been frequently employed in Raman spectral analyses for disease diagnosis such as AD, cancer, infectious disease, *etc.*.^22–25^ High accuracy in diagnosis is enabled by machine learning models including support vector machine (SVM),^26^ random forest classifier^27^ and neural networks.^28^ Besides achieving outstanding performance in classification, machine learning can also interpret the correlation between Raman modes and diseases by providing spectral feature importance map.^29, 30^ Such interpretability of machine learning can lead to key insights into the potential disease biomarkers by correlating spectral feature importance map with the signature molecular Raman spectra. However, so far machine learning interpretation lacks quantitative correlation to molecular composition in the Raman analysis for biomedical systems.^22–25^

In this work, we employed machine learning classification and interpretation on Raman spectra of mouse brain slices and screened AD biomarkers. Our workflow is primarily composed of three-step procedures: first, we collected Raman spectra on mice brain slices with and without AD; then, we used machine learning to classify the collected Raman spectra on AD and non-AD brain slices; finally, we used linear SVM to interpret the spectral feature importance map which differentiates AD and non-AD spectra and discovered potential AD biomarkers (Figure 1). In our Raman measurements, we used a special noise reduction technique: contacting monolayer graphene with the brain slices. Compared to intrinsic Raman spectroscopy, our unique graphene-assisted Raman spectroscopy enhanced Raman signal-to-noise ratios (from 53.9 to 121.0) and improved machine learning classification performance (accuracy from 77% to 98%). By comparing the machine learning prediction accuracy on Raman spectra from different brain regions, our experiment revealed that certain brain regions, such as the cortex, are more informative in AD identification. In our machine learning interpretation, the spectral feature importance map is found to register well with the Raman signatures of known AD biomarkers including Aβ and tau proteins. We also located several other molecules that have high Raman spectral correlation with the spectral feature importance map, which have been verified in previous biochemical studies, indicating their potential as the biomarkers for AD diagnosis. Combining Raman sensing and machine learning analysis to enable biomarker identification on brain slices, our interpretable machine learning based framework provides a route for fundamental study of AD pathology and can greatly accelerate AD diagnosis and drug development. Our Raman-machine learning integrated method has the potential to be extended to study other diseases and can be applied to various tissues, biofluids, and human samples.

**Figure 1.**
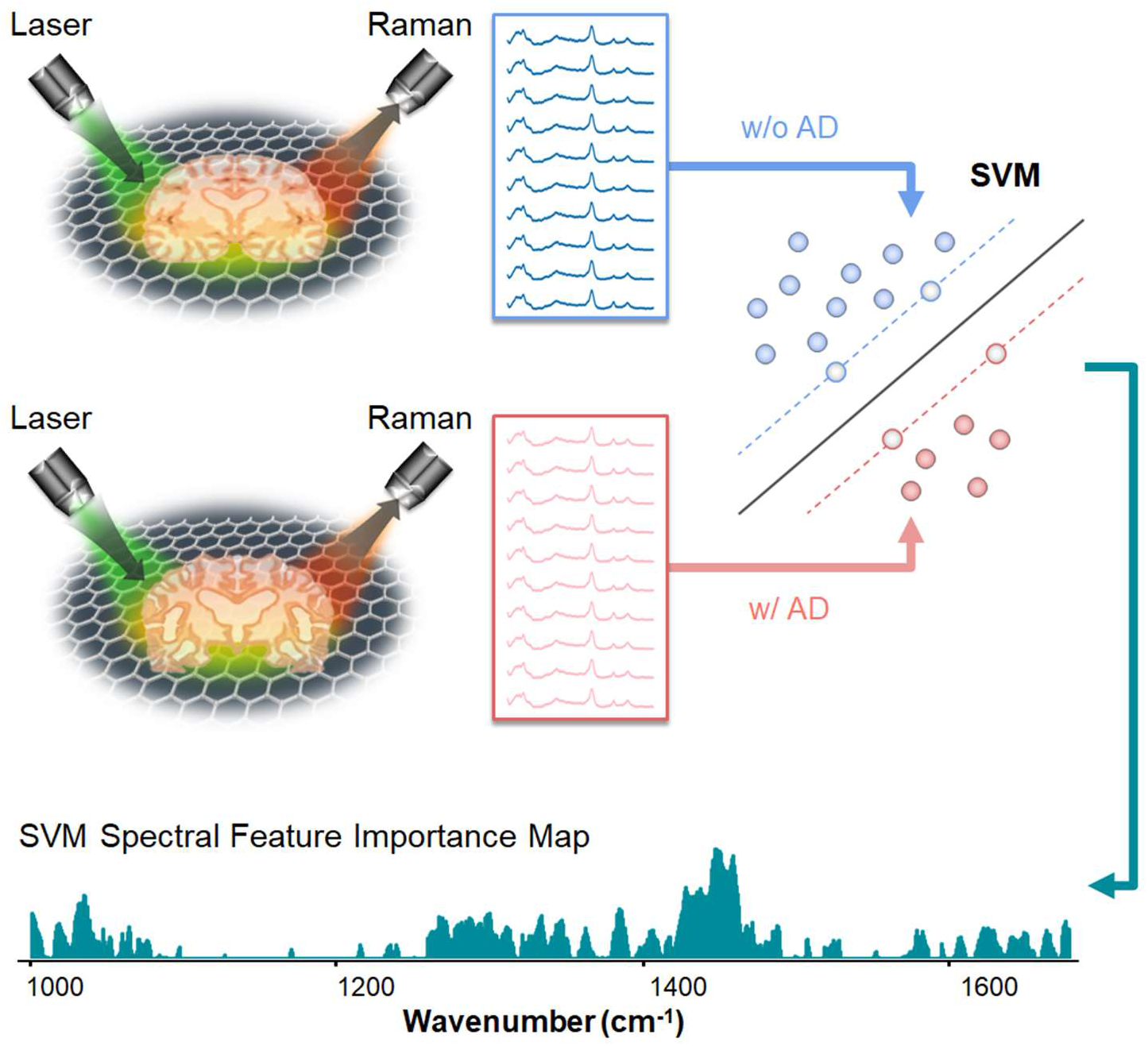
The overall workflow. Workflow of graphene-assisted Raman signals data collection, preprocessing, machine learning classification and interpretation. The machine learning classifier demonstrated is the linear SVM model to differentiate AD/non-AD Raman spectra.

## RESULTS AND DISCUSSION

### Improving the Signal-to-Noise Ratio Using Graphene-Assisted Raman Spectroscopy

We collected Raman spectra of brain slices harvested from AD transgenic mice and healthy mice. There are in total 351 spectra with AD and 376 spectra without AD in our dataset measured on three brain regions: cortex, hippocampus, and thalamus. During the Raman measurement, the brain slices were immersed in a neuroprotectant solution sealed between the silicon substrate and a fused quartz cover slide. For part of the measurements, we placed brain slices in direct contact with monolayer graphene which had been transferred onto the quartz cover slide (Figure S1).

Before feeding the Raman spectra into machine learning classifiers, we implemented the Savitzky-Golay filter^31^ for spectral smoothing and asymmetric least squares smoothing^32^ for baseline correction. Comparisons of raw Raman spectra before and after preprocessing are shown in Figure S2 and Figure S3. Figure 2a and 2b show examples of the preprocessed Raman spectra for AD and non-AD samples measured with and without graphene contact. As can be seen, there are major Raman peaks at 1038 cm^-1^,1088 cm^-1^, 1283 cm^-1^, 1312 cm^-1^, 1439 cm^-1^, 1458 cm^-1^ (Figure 2a and 2b). The graphene G band is at 1589 cm^-1^, noted as G (Figure 2a, Figure S4). The other Raman peaks are contributed by Aβ and tau proteins, and major molecular components in the brain (Table 1). For example, the 1283 cm^-1^ mode is contributed by the CH_2_ bending mode of oligomeric tau, actin, myelin basic protein, phosphatidylcholine, and triolein molecules. The Raman mode at 1458 cm^-1^ is contributed by the C – C stretching mode and CH_2_ bending mode of oligomeric Aβ, oligomeric tau, actin, glycogen, lactate, phosphatidylcholine, and triolein molecules.

**Figure 2.**
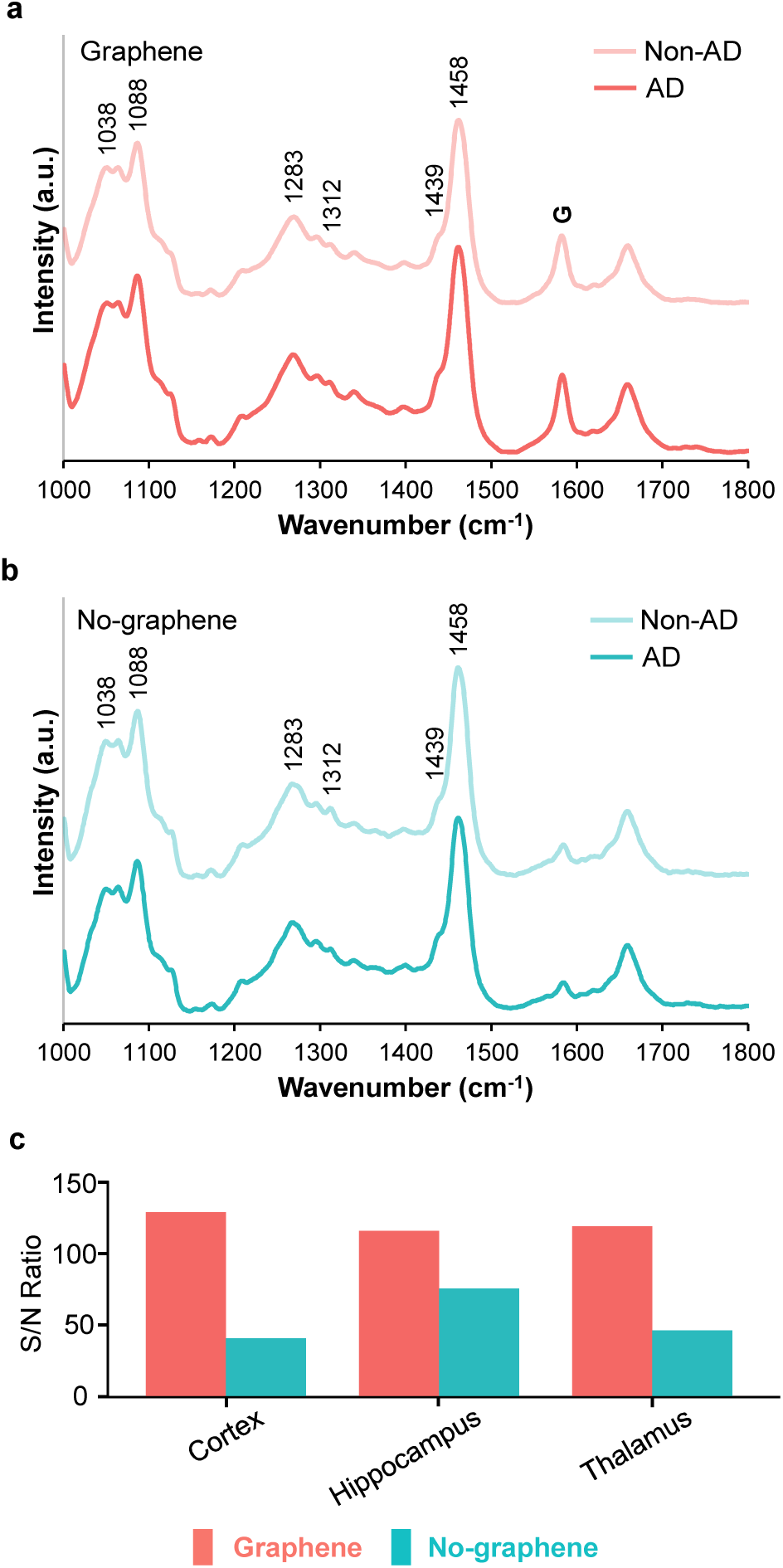
Raman spectra on brain slices. (a) Preprocessed Raman spectra in cortex region with and without AD, with graphene. Graphene G-band at 1589 cm^-1^ is notated as “G”. (b) Preprocessed Raman spectra in cortex region with and without AD, without graphene. (c) S/N of every brain region, measured with and without graphene.

**Table 1.** Assignments of most important Raman bands. Raman vibration modes, Aβ and tau proteins, and 17 major composite molecules of the brain assigned to major Raman peaks.^6,33–56^

| Peak position (cm<sup>-1</sup>) | Band assignment | Tentative contribution |
| --- | --- | --- |
| 1038 | (C – O)<sup>v</sup> | Oligomeric Aβ, Fibril Aβ; Actin, Glycogen, Myelin basic protein |
| 1063 | (C – C)<sup>v</sup> | Oligomeric Tau; Actin, Myelin basic protein, Phosphatidylcholine, Triolein |
| 1088 | (C – O – C)<sup>v</sup> | Aspartate amino, Ubiquitin |
| 1270 | Amide III |  |
| 1283 | (CH<sub>2</sub>)<sup>y</sup> | Oligomeric Tau; Actin, Myelin basic protein, Phosphatidylcholine, Triolein |
| 1297 | (CH<sub>2</sub>)<sup>y</sup> | Oligomeric Aβ; Actin |
| 1312 | (CH<sub>2</sub>)<sup>t</sup> | Monomeric Tau, Fibril Tau; Actin, Glycogen, Myelin basic protein, Phosphatidylcholine, Triolein |
| 1439 | (CH<sub>2</sub>)<sup>y</sup> | Monomeric Aβ, Oligomeric Tau, Fibril Tau; Aspartate Amino, Lactate, Cholesterol, Myelin basic protein |
| 1458 | (C – C)<sup>v</sup>,<br>(CH<sub>2</sub>)<sup>y</sup> | Oligomeric Aβ, Oligomeric Tau; Actin, Glycogen, Lactate, Phosphatidylcholine, Triolein |
| <sup>v</sup>stretching, <sup>y</sup>bending, <sup>t</sup>twisting |  |  |

We calculated the signal-to-noise ratios (S/N) for spectra with and without graphene in three brain regions. The S/N are shown in Figure 2c where red bars correspond to graphene-assisted spectra and blue bars represent the no-graphene results. It is clear that the graphene-assisted spectra exhibit much higher S/N than the no-graphene spectra for all the three brain regions (S/N average of 121.0 for graphene-assisted spectra compared to 53.9 for no-graphene spectra). Overall, we observed that when brain slices were placed in contact with graphene, the Raman spectra exhibited less noise when compared to the measurements without graphene. Our prior work, along with others, has shown that graphene can reduce the noise of Raman spectra, enhance Raman signals, and quench fluorescence for organic and biomolecules.^57–60^ Here, the reduced noise can be attributed to the above factors, as well as the high heat conductivity of graphene which can reduce the laser heating effect during Raman measurements of brain slices.^61, 62^ Therefore, we used graphene-assisted Raman spectra for further investigation described in the following sections.

### Machine Learning Classification

In order to classify AD and non-AD spectra, we used the graphene-assisted spectra and applied different algorithms including linear SVM,^26^ random forest,^27^ XGBoost,^63^ CatBoost.^64^ The common metrics for machine learning including classification accuracy, area under the receiver operating characteristic curve (AUC), sensitivity, and specificity for graphene-assisted Raman spectra are shown in Figure 3. Although the Raman spectra for AD and non-AD brain slices are visually similar (Figure 2a), machine learning classification can capture minor differences and distinguish the two classes with high accuracy. It can be seen from Figure 3 that the cortex region rendered the accuracy over 93% for every classifier. Compared to the hippocampus and thalamus, the cortex region exhibited better accuracy, AUC, sensitivity, and specificity for all classifiers. Thus, we can infer that Raman fingerprints of AD-relevant biomarkers are better captured in the cortex region with graphene assistance.^65^ It should be noted that our results do not indicate that the roles of hippocampus and thalamus in AD should be ignored since our observations only indicate that graphene-assisted Raman signal is more sensitive to AD-relevant molecular components in the cortex region compared to the other two regions. We also performed the same machine learning classification for Raman spectra measured without graphene, and the results are shown in Figure S5 and Table S1.

**Figure 3.**
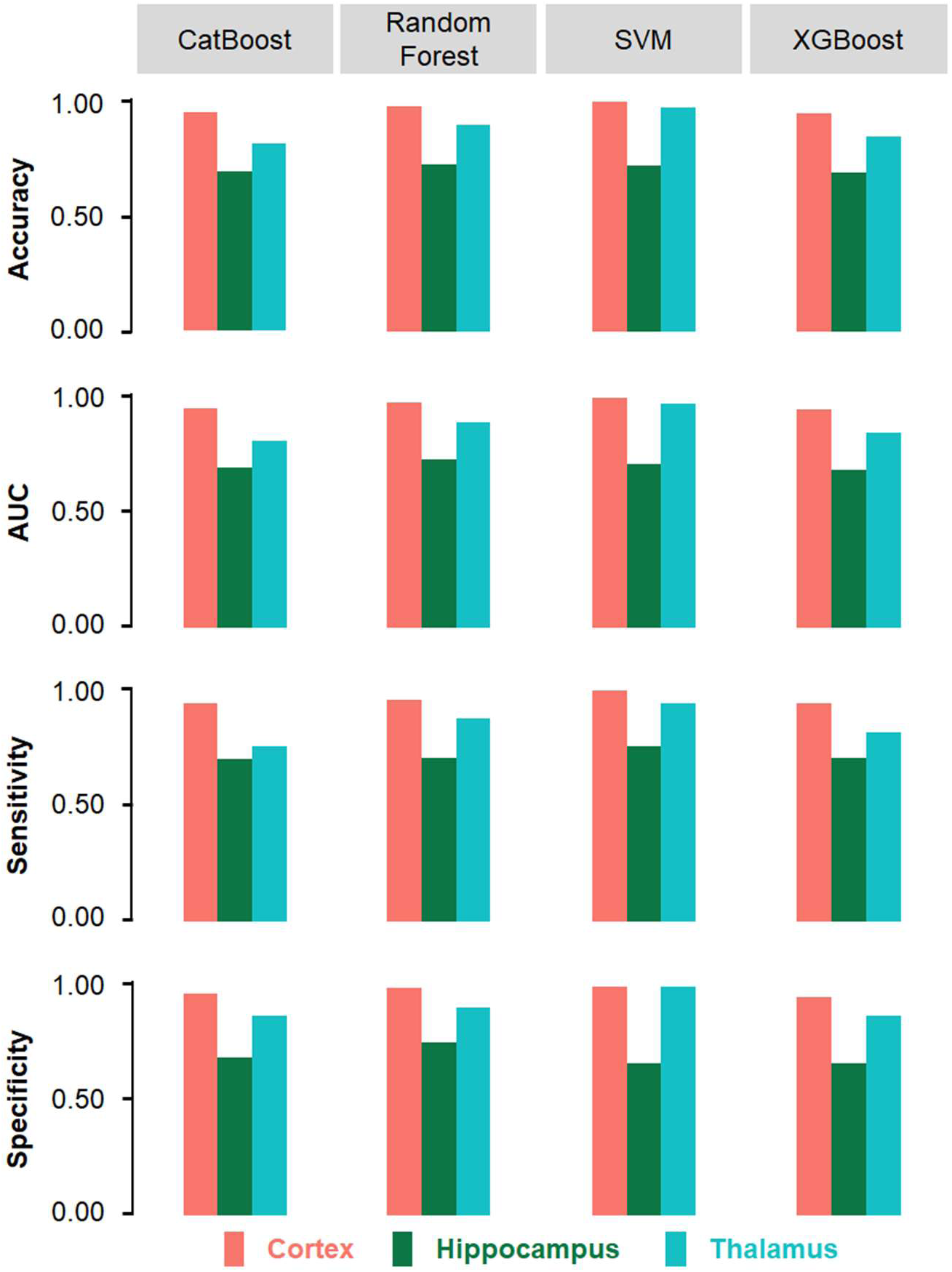
Machine learning classification summary on graphene-assisted Raman spectra. Accuracy, AUC, sensitivity, and specificity of graphene-assisted Raman spectra from the cortex, hippocampus, and thalamus regions.

As the graphene-assisted Raman signals from the cortex region yield better results in the machine learning classification, we visualized the data distribution using t-distributed stochastic neighbor embedding (t-SNE) plots, a non-linear dimensionality reduction technique, for selecting appropriate interpretable machine learning classifiers.^66^ As shown in Figure 4a, in the cortex region, the graphene-assisted Raman data can be well separated by a linear decision boundary while the Raman data measured without graphene are apparently not linearly separable (Figure 4b). Meanwhile, the machine learning classification accuracy using graphene-assisted Raman data reaches as high as 98% using linear SVM; however, the accuracy is at most 77% among the four classifiers using the no-graphene data (Figure 4c). This, again, shows the high quality of our graphene-assisted Raman data, which are more suitable for feature importance matching and interpretation (t-SNE plots from other brain regions are in Figure S6). A linear classifier with high accuracy is preferable to perform this interpretation task, since it is simple yet sufficient to fit the linearly-separated data without introducing much model complexity, making its spectral feature map more straightforward for interpretation. Thus, we chose linear SVM from the series of machine learning models experimented for further investigation to determine candidate biomarkers in association with AD.

**Figure 4.**
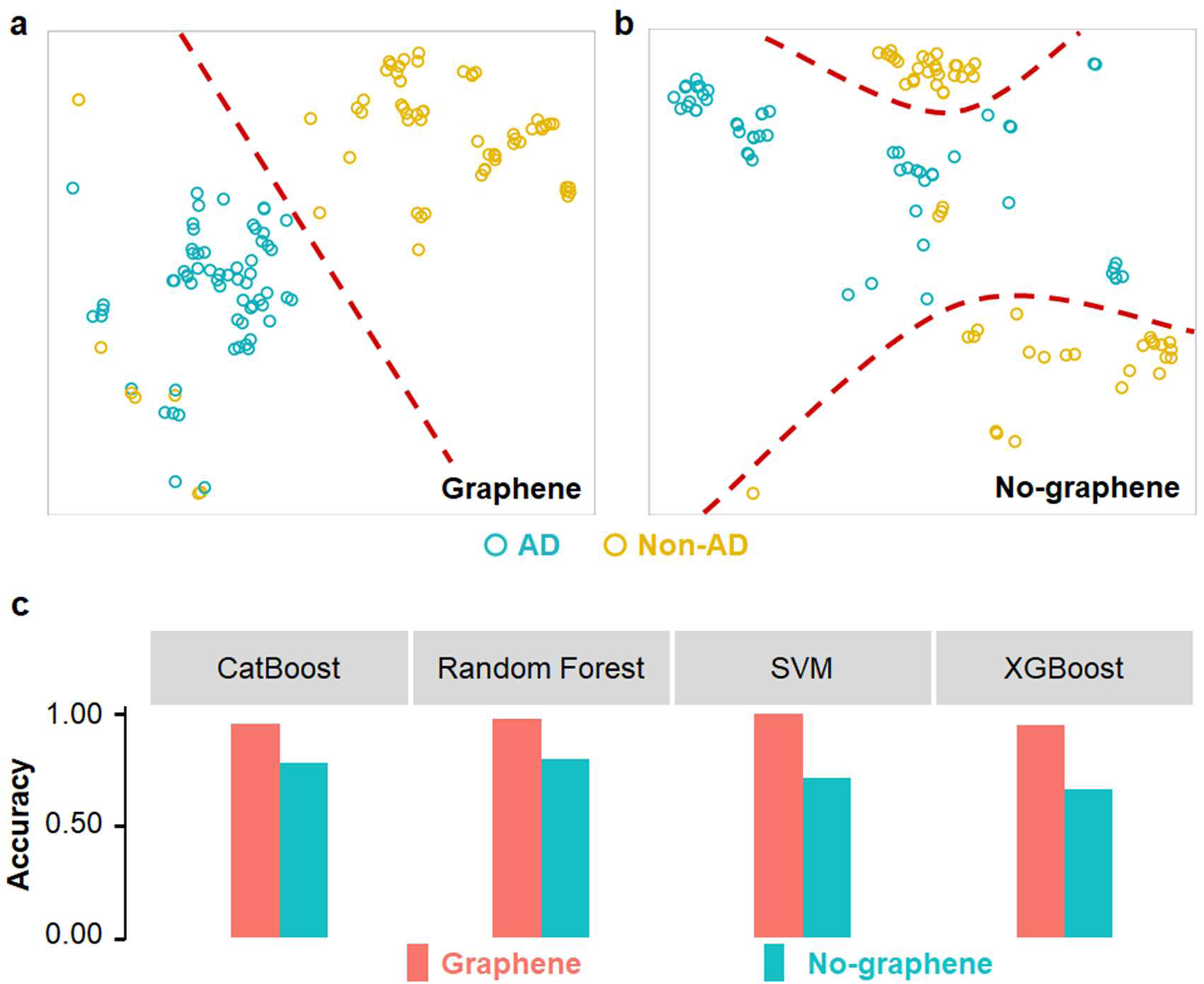
t-SNE plots for cortex region. (a) t-SNE plot of graphene-assisted spectra on cortex region. The red dashed line is the estimated linear decision boundary. (b) t-SNE plot of spectra without graphene on cortex region. The red dashed curves are estimated non-linear decision boundaries. (c) Comparison of classification accuracy between Raman spectra with and without graphene on cortex region.

### Machine Learning Interpretation

We further interpreted the machine learning classification results and studied the features learned, which can provide important information on AD biomarker molecules. Validated by high accuracy, AUC, sensitivity, and specificity (Figure 3), the features learned by SVM from the graphene-assisted data in the cortex region are considered reliable for interpretation. From the trained linear SVM model, we assigned a spectral mapping of Raman wavenumbers based on their importance in classifying AD/non-AD spectra. The extracted spectral feature importance map contains two sets of features: positive features and negative features (positive features shown in Figure 5, and complete features shown in Figure S7). Since the spectral feature importance map shows the importance of each Raman wavenumber in the AD/non-AD classification, it stresses the difference between AD and non-AD Raman spectra.

**Figure 5.**
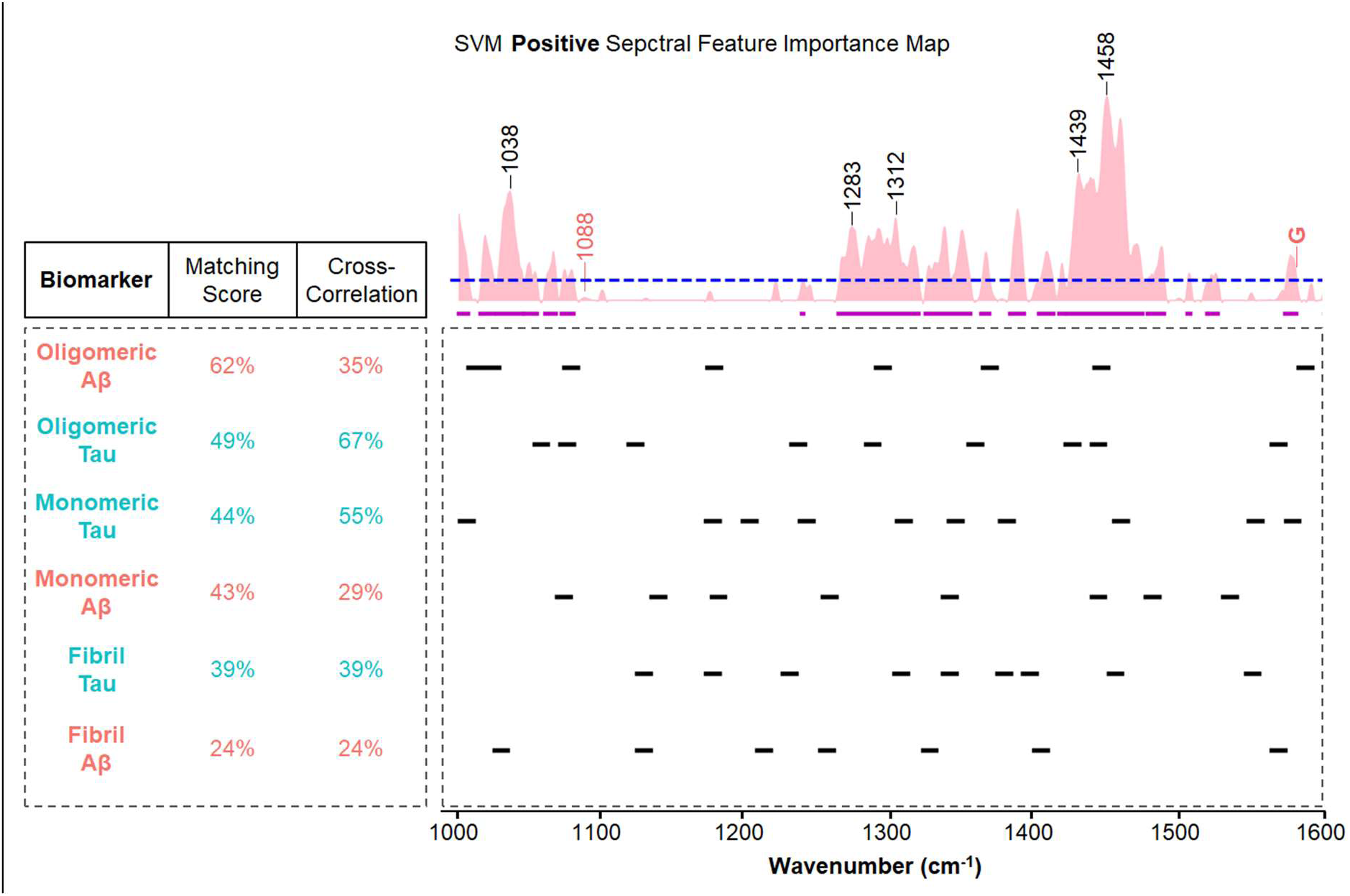
Feature importance matching for AD biomarkers. (Top) The positive spectral feature importance map found by SVM, where wavenumbers with high feature importance scores (*i.e.* high values along the vertical axis) are important for AD recognition. On the spectral feature importance map, the blue dashed line is the threshold of positive important feature ranges used for matching scores. The purple solid lines are important feature ranges above the threshold. Important peaks corresponding to signature Raman modes are labeled in black. Non-important peaks corresponding to signature Raman modes are differentiated in red. (Bottom left) The matching scores and cross-correlation coefficients between SVM feature importance map and previously reported 6 biomarkers.^51–53^ (Bottom right) The black solid lines below the spectral feature importance map are Raman signature peak ranges of 6 biomarkers with plus/minus 5 cm^-1^. The wavenumber ranges from 1000 to 1600 cm^-1^ .

As shown in Figure 5, the most important features from machine learning interpretation are 1038 cm^-1^, 1283 cm^-1^, 1458 cm^-1^, whose importance values are 0.087, 0.056, 0.15, respectively. All of these wavenumbers have Raman peaks for both AD and non-AD samples, but there is a slight difference: the AD samples exhibit about 10.5 ± 7.4%, 12.5 ± 3.5%, and 12.6 ± 1.4% stronger intensity than the non-AD samples, respectively, as shown in Figure 2a, which matches our analysis results that these wavenumbers possess important positive spectral features. The Raman mode at 1038 cm^-1^ may correspond to C – O stretching mode in oligomeric Aβ and fibril Aβ proteins. It may also be contributed by actin, glycogen, and myelin basic protein molecules. The 1283 cm^-1^ may correspond to CH_2_ bending mode in oligomeric tau protein. It may also be contributed by actin, myelin basic protein, phosphatidylcholine, and triolein molecules. The 1458 cm^-1^ may correspond to C – C stretching mode and CH_2_ bending mode in oligomeric Aβ and oligomeric tau proteins. It may also be contributed by actin, phosphatidylcholine, and triolein molecules. The high spectral feature importance in those wavenumbers suggests that the biological molecules mentioned here are potentially related to the diagnosis of AD. On the other hand, if the Raman peaks identified in both AD and non-AD samples do not exhibit significant spectral difference, the spectral feature importance does not necessarily show peaks in that wavenumber. For example, the graphene G band at 1589 cm^-1^ in both AD and non-AD spectra does not appear as an important wavenumber in Figure 5 since graphene is deployed in the same way in both AD and non-AD samples. Also, the Raman peak around 1088 cm^-1^ for both AD and non-AD samples (Figure 2a), which is potentially related to C – O – C stretching mode for aspartate amino and ubiquitin molecules, is not important according to our spectral feature importance, since the aspartate amino and ubiquitin molecule is not closely associated with AD. The interpretability of machine learning we demonstrated here presents a tremendous advantage of machine learning interpretation and enables the discovery of biomarkers that have very small quantities in the diseased samples, potentially allowing for early-stage diagnosis and understanding of disease pathology.

To better understand our spectral feature importance map and its relationship with molecular components, we developed two metrics for two application scenarios: Pearson cross-correlation coefficient based algorithm (if Raman spectra of biomarkers, including peak frequencies and intensities, are available), and matching score based on spectral overlap between important feature ranges and biomarker Raman peaks (if only the peak frequencies of biomarkers are known, and peak intensities are unavailable). Note that the former metric is relatively informative since it considers all spectral features including frequencies and intensities. On the other hand, the latter metric based only on peak frequency is relatively robust since peak frequency is stable compared with other spectral features such as intensity, which depends on the measurement conditions such as laser wavelength, laser power, and substrate. For validation, here we used both metrics to cross-check our interpretation results. We first examined several major AD biomarker proteins using both metrics. From the results shown in Figure 5, it is obvious that the known AD biomarkers such as oligomeric tau (with Raman peaks at 1063, 1283, 1439, 1458 cm^-1^, *etc.*) and oligomeric Aβ (with Raman peaks at 1038, 1297, 1458 cm^-1^, *etc.*) have considerable cross-correlation coefficients (Metric #1) and matching scores (Metric #2). To be more specific, oligomeric tau has a cross-correlation coefficient of 67% and a matching score of 49% and oligomeric Aβ has a cross-correlation coefficient of 35% and a matching score of 62%, while other uncorrelated molecules such as tropomyosin and hemoglobin beta only have cross-correlation coefficients and matching scores no more than 13%. Both of our metrics correctly identified the significance of Aβ and, meanwhile, recognized the role of tau in AD brain slices. This demonstrates the validity of the Pearson cross-correlation coefficient and matching score metrics that we developed.

Using the above two metrics, we can further screen more molecules that are potentially correlated to AD. We applied the cross-correlation coefficient and matching score to 17 major composite molecules of the brain (Table S2). The spectra of the component molecules with cross-correlation coefficients above 65% are shown in Figure 6. Triolein, phosphatidylcholine, and actin have the highest cross-correlation coefficients to spectral feature importance map of 72%, 71%, 69%, respectively. Consistent with the cross-correlation coefficient metric, the matching scores of these three molecules are also the highest among the 17 composite molecules. As clearly seen in Figure 6, the signature Raman peaks of triolein molecule, phosphatidylcholine molecule, and actin molecule including 1283 cm^-1^ (CH_2_ bending mode), 1312 cm^-1^ (CH_2_ twisting mode), and 1458 cm^-1^ (C – C stretching mode and CH_2_ bending mode) match well with both our spectral feature importance map (Figure 6) and Raman peak analysis (Figure 2a and Table 1). This suggests that triolein, phosphatidylcholine, and actin may be associated with AD, which have been suggested by prior reports of biochemical and physiological studies.^67–69^ For example, Bamburg *et al.* found an increase of actin in the AD brain compared to the normal brain.^67^ Johnson *et al.* suggested that triglycerides, including triolein, can also lead to cognitive impairments., where they reported concentration levels of 46.49 mg/dl for AD animals and 35.01 mg/dl for healthy animals.^70^ Banks *et al.* also showed that the decrease of triglycerides improves both learning and memory capabilities.^68^ Additionally, phosphatidylcholine is found to be significantly lower in AD patients,^71^ where high levels of phosphatidylcholine usually reduce the progression of dementia.^72^ However, it is worth noting that the AD pathogenesis is different in mice and human brains in terms of the phosphatidylcholine level. According to Chan *et al.*, phosphatidylcholine levels are lower in the AD human brain while higher in the mouse forebrain (18 Mol% for AD animals and 16 Mol% for healthy animals).^69^

**Figure 6.**
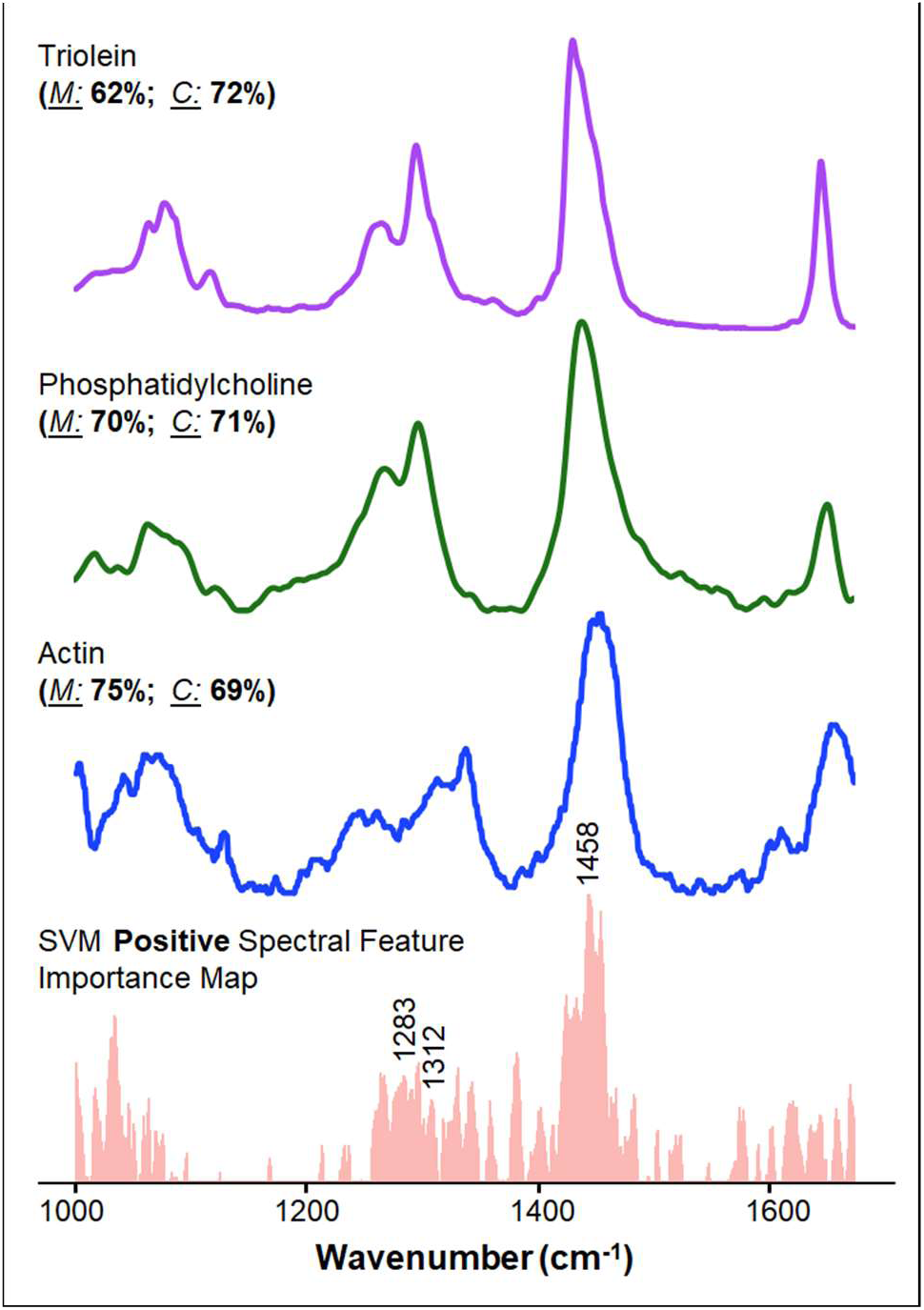
Other potential AD biomarkers identified. Biomolecular component spectra with the top 3 cross-correlation coefficients to the spectral feature importance map: triolein, phosphatidylcholine and actin.^39, 41, 49^ *<u>M</u>* refers to the matching score, and *<u>C</u>* refers to the cross-correlation coefficient.

In addition to the positive features shown in Figures 5 and 6, the SVM classifier also found negative features in the range 1100-1250 cm^-1^, *i.e.*, the representative Raman spectral range for non-AD brain slices (Figure S7). This means that biomolecules with Raman peaks in this wavenumber range tend to be negatively correlated with AD (*i.e.* they likely reduce in amount or disappear with AD). These negative features correspond to cytochrome (with Raman peak at 1139 cm^-1^) and glycogen (with Raman peak at 1237 cm^-1^) in both matching metrics, suggesting that cytochrome and glycogen are negatively correlated to AD, again consistent with prior biochemical experiments.^73–76^ In earlier studies, glycogen has been purposed to have beneficial effects for cognition,^73^ whose negative correlation has been confirmed by Bass *et al.*^74^ Furthermore, a decreased level of cytochrome has also been found in AD brain slices. For example, cytochrome c oxidase is found decreased in AD as reported by Parker *et al.* and Cardoso *et al.*.^75, 76^

We note that our approach is able to screen potential biomarkers, rather than to accurately pinpoint all the biomarkers which requires further biochemical verification. The accuracy of our matching score and cross-correlation coefficient metrics depends on the quality and range of molecular Raman data available in the literature. For instance, the matching score and cross-correlation coefficient values in Figure 5 and Figure 6 are not directly comparable, because the Raman signal range used was 1000 – 1600 cm^-1^ for results shown in Figure 5, and the Raman range used was 1000 – 1670 cm^-1^ for results shown in Figure 6. The smaller Raman range for biomarker analysis (Figure 5) was due to the limited Raman spectral range reported for Aβ and tau proteins in the literature.^53^ Although not reported before, Aβ and tau proteins might have many important Raman modes between 1600 – 1670 cm^-1^, such as amide I mode at 1665 cm^-1^ for oligomer Aβ protein and amide I mode at 1654 cm^-1^ for fibril Aβ protein,^51, 52^ which could affect the matching scores and cross-correlation coefficients of Aβ and tau in our analysis. We further note that our biomarker screening method in principle is not limited by the number of biomarkers. To identify more potential biomarkers, we simply input the Raman spectra of other biomolecules within the same spectral range (*e.g.* from a Raman spectra database) in our component matching with spectral feature importance map.

## CONCLUSION

In summary, we measured and analyzed Raman spectra on mice brain slices with and without AD and used machine learning to classify AD/non-AD spectra in order to screen biomarkers through the interpretation of spectral feature importance. Raman spectra, with their multiple and narrow Raman peaks, contain rich molecular information and potentially provide an ideal dataset for machine learning analysis. Our graphene-assisted Raman measurements demonstrated further enhanced S/N, and thus effectively improved the performance of machine learning classification to achieve a high accuracy of 98%. To further interpret the molecular information in the Raman spectra, we obtained calculated spectral feature importance map based on our machine learning classifier, and developed two new metrics, including cross-correlation coefficient and matching score, to identify molecules that are relevant to AD. Our interpretable machine learning based framework recognized a series of known AD biomarkers such as oligomeric tau and oligomeric Aβ that have a considerable correlation to AD. Our model also identified three molecules (triolein, phosphatidylcholine, and actin) that are positively correlated to AD, and two molecules (cytochrome and glycogen) that are negatively correlated to AD. Our work offers a rapid approach to detect AD and to screen AD biomarker molecules, thus is promising to greatly accelerate the study of AD in terms of diagnosis and treatment. Our approach integrating graphene-assisted Raman spectroscopy and interpretable machine learning can also be widely applied to study various other diseases and to a wide range of biological samples including tissues and biofluids.

## METHODS

### Animals and brain slice preparation

Animal investigation procedures were conducted in accordance with institutional and NIH guidelines. The animals were housed with *ad libitum* access to food and water in a room with a 12-hr light and dark cycle in the animal facility. We utilized the previously reported 5XFAD mouse model expressing APP^Swedish/Florida/London^ and PSEN1^M146L/ L286V^ mutations,^77, 78^ which recapitulates the features of Alzheimer’s β-amyloid pathology in animal brains. 5XFAD mice of different ages and non-transgenic (non-AD) control mice were investigated. All animal study procedures were approved by MGH IACUC (Protocol #: 2011N000022).

Brain slices were prepared following previously reported methods.^79, 80^ Particularly, animals were anesthetized with isoflurane and then decapitated. Isolated brains were longitudinally bisected, and hemispheres were separated and incubated in 4% paraformaldehyde-containing 0.1 M phosphate-buffered saline (PBS) at 4 °C for 48 hrs, followed by incubation in 30% sucrose solution in 0.1 M PBS. Next, brains were snap frozen from a dry ice-cooled block on a sliding microtome (Leica SM 2010R), and sectioned in 40 μm thickness. The free-floating brain sections were stored at −20°C in a cryoprotective buffer containing 28% ethylene glycol, 23% glycol, and 0.05 M PBS, until subsequent analysis by Raman testing.

### Raman measurement

The Raman measurement was performed on the Horiba LabRam system with a 50× objective. The laser power on the sample was controlled below 0.4 mW to avoid potential laser bleaching. The excitation laser wavelength was 532.5 nm. All the measurements were performed on mice brain slices in the neuroprotectant solution that was sealed between a quartz cover slide and the silicon substrate. In the case with graphene in contact, the quartz slide has monolayer graphene transferred on the surface. In each Raman measurement, we carried out 3 accumulations, thus each spectrum is in effect averaged 3 times.

### Graphene synthesis and transfer

The graphene layers were synthesized by chemical vapor deposition method. For the graphene growth, Cu foil was first placed in a quartz tube furnace and annealed at 1065°C for 1hr under 60 sccm H_2_ and 940 sccm Ar at atmospheric pressure. For the growth, the furnace was brought down to 1000°C and the gas flow rates were updated to 36 sccm H_2_ and 2204 sccm Ar. Then 0.6 sccm CH_4_ was introduced for 1hr. After 1hr, the CH_4_ was turned off and the Cu foil was rapidly cooled by removing it from the furnace area.

For the transfer, a supporting PMMA layer was spin-coated on top of the graphene/Cu stack. Then the Cu was etched away using commercial Cu etchant. After the Cu has etched, the PMMA/graphene was put into three separate water baths for several hours, before being transferred to the desired final substrate. After the sample has dried, it was placed into acetone overnight in order to remove the PMMA and then rinsed with isopropyl alcohol and blow dried.

### Data preprocessing and calculation of signal-to-noise ratio

After obtaining the raw data, it is essential to apply preprocessing methods to reduce the effect of noise and background on classifiers. For each spectrum, we applied Savitzky-Golay filter to reduce the spectral noise.^31^ We removed the background using baseline correction with asymmetric least squares smoothing.^32^ To calculate S/N, we selected peaks of interest at 1038 cm^-1^, 1088 cm^-1^, 1283 cm^-1^, 1458 cm^-1^ and 1649 cm^-1^. Then, we divided the average intensity of each peak by the standard deviation of the peak intensity across different AD spectra from the same brain region to calculate the S/N for a single peak. Finally, we averaged the S/N of all peaks of interest to get the S/N of spectra of the area. Notice that since we used the same preprocessing for all (both graphene and no-graphene) spectra, comparison between S/N of graphene and no-graphene spectra are not affected.

### Classifiers architecture and feature importance

As shown in Figure 3, multiple classifiers were used in the experiment including linear SVM, random forest, XGBoost, and CatBoost. Classification experiments were implemented using stratified 5-fold cross-validation to preserve the same percentage of samples for each class to improve robustness. For experiments with a relatively small sample set, SVM is usually an efficient and reliable option, for it is designed to find the optimal decision boundary represented by a hyperplane that maximizes the margin of separation between different classes.^24^ In our binary classification, we used linear SVM to find the optimal linear decision boundary between the two classes. The sign of a feature weight obtained from the Linear SVM classifier represents that feature’s direction to predict class.^81, 82^ Hence, the feature weights can be intuitively interpreted as spectral feature importance map shown in Figure 5 (positive only), while the positive (negative) features correspond to Raman signals more represented in AD (non-AD) samples. We used scikit-learn package to implement linear SVM and extract spectral feature importance map.^83^

### Component Cross-correlation Coefficient for Raman

*Pearson correlation coefficient* measures the linear correlation between two variables.^84^ The method is a standard measure of similarity between two Raman spectra.^85^ Here, we used the cross-correlation coefficient to measure the levels of correlation between the machine-learning derived feature map and the Raman spectra of 17 commonly known components in the brain from literature (Table S2).^33–48^ We modified the Pearson cross-correlation coefficient method and excluded negatively correlated trends as shown as follows:

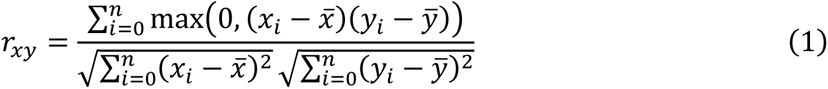

Considering the range difference of each spectrum, we normalized the components’ spectra and the spectral feature importance map so that they were within the same interval (1000 – 1600 cm^-1^) and have the same dimension. Then we used *Equation 1* to calculate the cross-correlation coefficient. This results in a coefficient that ranges from [0,1] where *r=0* means no correlation and *r=1* means perfect correlation.

### Matching Score Between Machine-learning derived Feature Map and Biomolecule Raman Spectra

Raman peak intensities vary with a number of factors including excitation wavelength and substrate. To avoid the influence of Raman peak intensity variation, we developed another metric, the *Matching Score,* for measuring the correlation between biomarkers and distinguishable patterns learned from machine learning models to some extent. The metric is designed as a ratio, with the numerator as the sum of important feature ranges of the extracted feature map of a particular class (*i.e.* either AD or No AD) that overlap with Raman spectra peak ranges of a certain biomarker, the sum of the aforementioned biomarker peak ranges is used as the denominator. Significant Raman peak ranges of 6 biomarkers (Fibril tau, Fibril Aβ, Oligomeric tau, Oligomeric Aβ, Monomeric tau and Monomeric Aβ) that are commonly known to be present in brain slides are gathered from literature.^51–53^ As demonstrated in Figure 5, Raman peak ranges are constructed by granting a shift of 5 wavenumbers for each biomarker representing important feature regions, the sum of which is utilized as the denominator for further calculations. Similarly, pinpointing the important feature ranges for the feature map as numerator is also desired. Rather than extracting peaks from feature map, we applied a 40% percentile threshold as the cutoff, regions above which we considered as significant and used the sum of their intersection with the denominator as the numerator. Matching scores of all six biomarkers and the positive feature map extracted from Linear SVM training are presented in Figure 5, indicating that results from our *Matching Score* approach are consistent with biomedical findings using other methods.

## ASSOCIATED CONTENT

### Supporting Information

The Supporting Information is available free of charge at https://pubs.acs.org/doi/xxx.

## AUTHOR INFORMATION

### Authors

**Z. Wang, K. Zhang, L. Ding and J. Ranasinghe** -Department of Electrical Engineering, The Pennsylvania State University, University Park, 16802, USA

**J. Ye -**College of Information Sciences and Technology, The Pennsylvania State University, University Park, 16802, USA

**T. G-Nakajima, M. Terrones -**Department of Physics, The Pennsylvania State University, University Park, 16802, USA

**Y. Xue -**Department of Electrical and Computer Engineering, Johns Hopkins University, Baltimore, 21218, USA

**S. Sharma, I. Biase -**Department of Computer Science, The Pennsylvania State University, University Park, 16802, USA

**S. H. Choi, R. E. Tanzi -**Genetics and Aging Research Unit, McCance Center for Brain Health, MassGeneral Institute for Neurodegenerative Disease Department of Neurology, Massachusetts General Hospital, Harvard Medical School, 114 16th Street, Charlestown, MA, 02129, USA

**C. Ran -**Martinos Center for Biomedical Imaging, Massachusetts General Hospital, Harvard Medical school, 13th Street, Building149, Charlestown, MA, 02129, USA

### Author Contributions

Z.W., J.Y., L.D. and J.R. performed experiments and analyzed data. T.G-N. and M.T. synthesized graphene. Y.X., S.S. and I.B. helped with data analysis. S.H.C., C.R., R.E.T., and C.Z. engaged in animal studies and interpreted results of brain slices. S.X.H., C.Z., and S.H. designed and supervised the work. Z.W., J.Y. and K.Z. wrote the manuscript draft and prepared the figures. All authors reviewed and revised the manuscript to its final state.

### Funding Sources

The authors thank the Imaging Facility of Materials Characterization Lab at Penn State for instrument use. C.Z. and S.H. acknowledge the support from the National Institutes of Health under grant number R56AG062208. L.D. and S.H. acknowledge the support from Johnson & Johnson Inc. for the STEM2D Scholar’s Award. S.H. acknowledges the support from the National Science Foundation under grant number ECCS-1943895. C.Z. acknowledges the support from the National Institutes of Health under grant number R01AG055784. C.Z. and R.E.T. acknowledge the support from the Cure Alzheimer’s Fund.

## Table of Contents

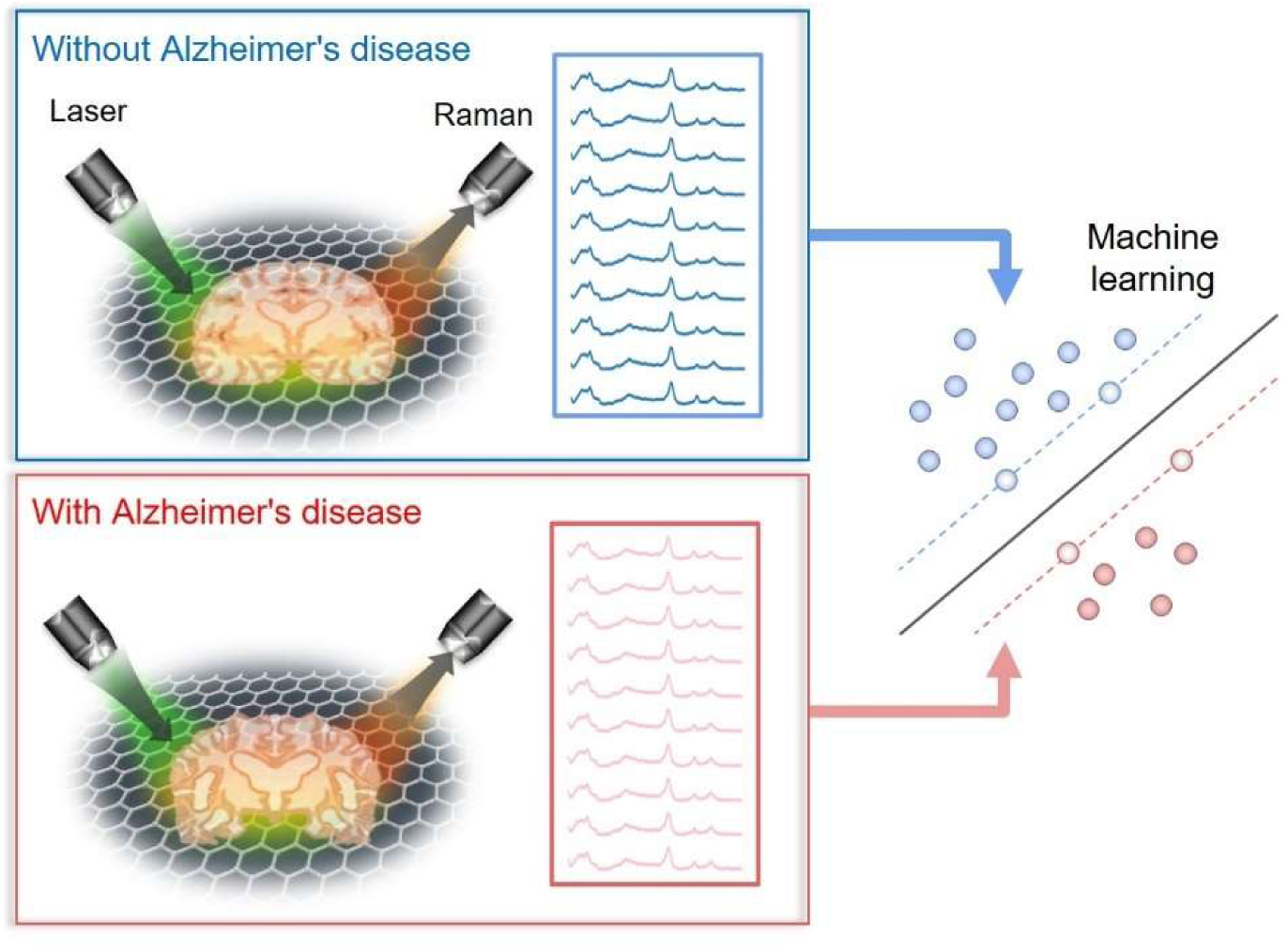

## Notes

### Competing Interest Statement

The authors have declared no competing interest.

### Summary of Updates

New experiment and data visualization.

