## Supporting Information for "Rapid Biomarker Screening of Alzheimer’s Disease by Interpretable Machine Learning and Graphene-Assisted Raman Spectroscopy"

**This PDF file includes:**

Figure S1 to S7
Table S1 to S2

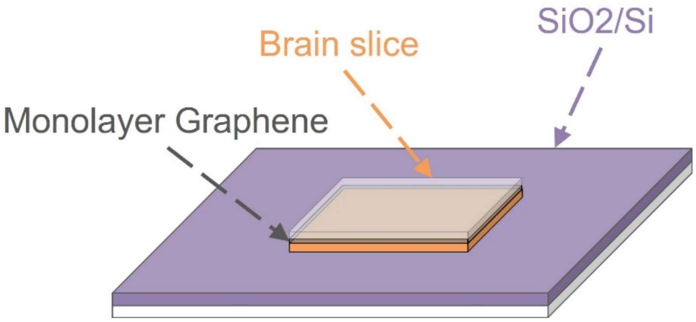


Figure S1. **Illustration of graphene-assisted Raman spectroscopy measurement.** Brain slices are in direct contact with monolayer graphene which was transferred on the quartz cover slide.

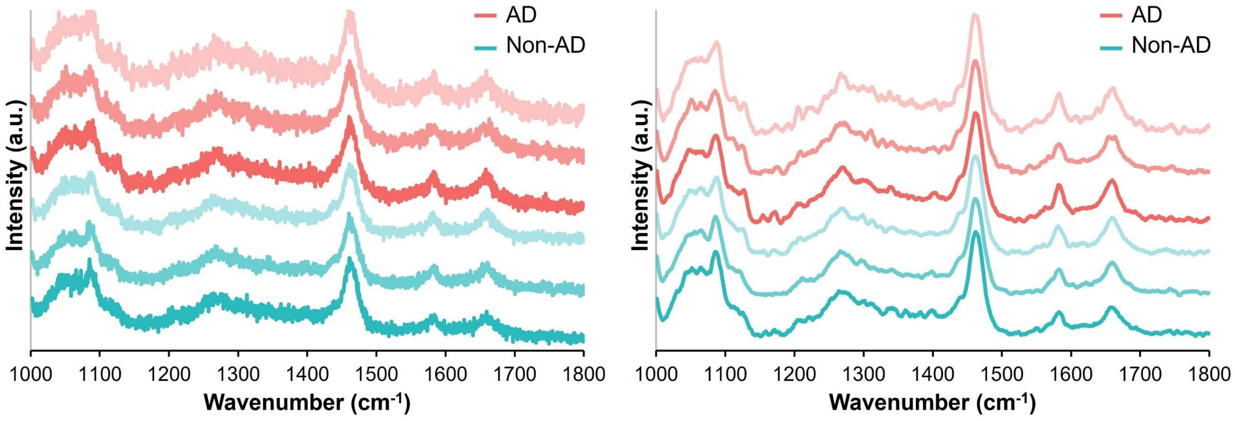


Figure S2. **Raman spectra on brain slices before and after preprocessing.** (Left) Graphene-assisted Raman spectra of AD and non-AD sample in cortex region before preprocessing. (Right) Graphene-assisted Raman spectra of AD and non-AD sample in cortex region after preprocessing.

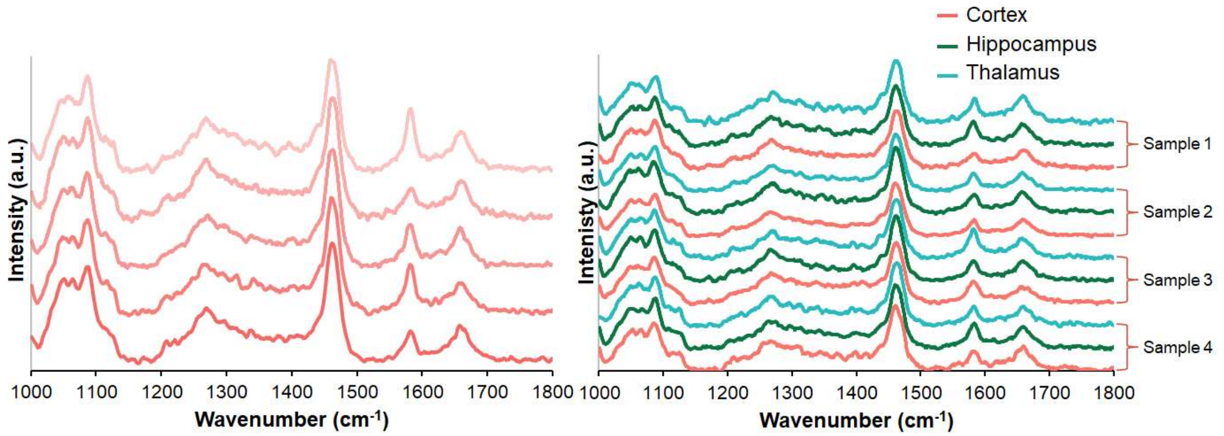


Figure S3. **Raman spectra on brain slices after preprocessing.** (Left) Graphene-assisted Raman spectra of preprocessed non-AD sample in hippocampus collected at different times. (Right) Graphene-assisted Raman spectra of preprocessed sample across different animals and brain slices.

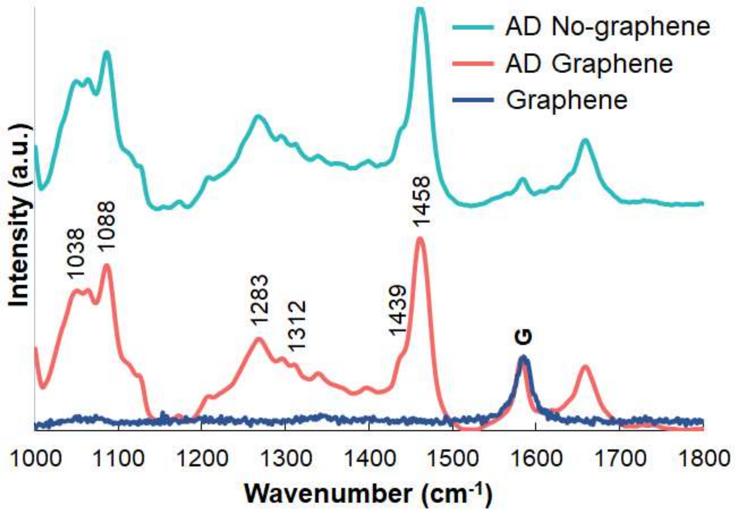


Figure S4. **Comparison between Raman spectra in cortex region and graphene peaks.** Preprocessed Raman spectra in cortex region with AD, with and without graphene. The Raman spectrum of pristine graphene on silicon. Graphene G-band at 1589 cm-1 is notated as “G”.

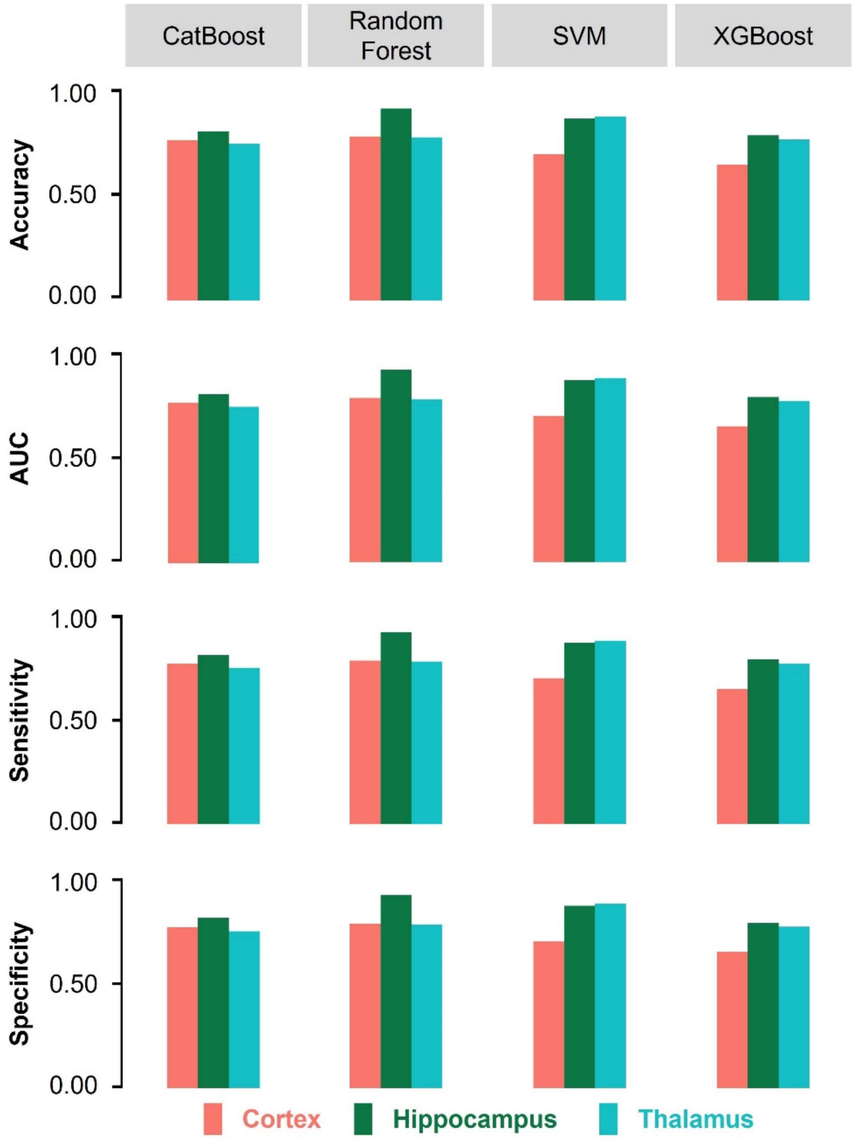


Figure S5. **Machine learning classification summary on no-graphene Raman spectra.** Accuracy, AUC, sensitivity, and specificity of Raman spectra without graphene from the cortex, hippocampus, and thalamus regions.

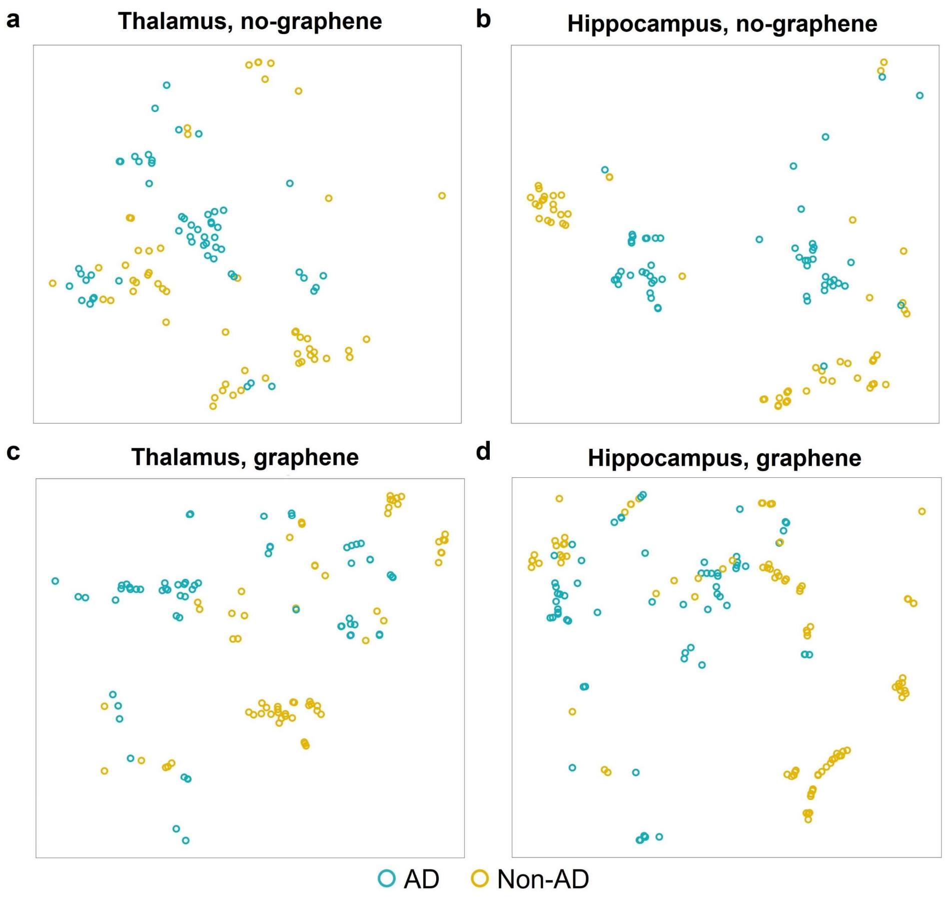


Figure S6. **t-SNE plots for thalamus and hippocampus regions.** (a) t-SNE plot of no-graphene spectra on thalamus region. (b) t-SNE plot of no-graphene spectra on hippocampus region. (c) t-SNE plot of graphene-assisted spectra on thalamus region. (d) t-SNE plot of graphene-assisted spectra on hippocampus region.

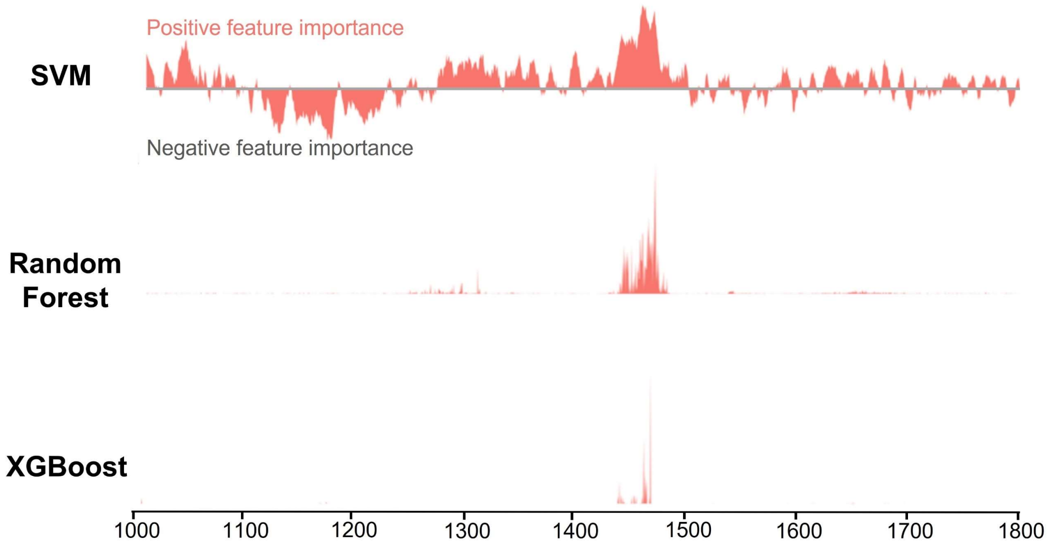


Figure S7. **Feature importance maps by linear SVM, random forest and XGBoost.** Notice that SVM is with both positive and negative feature importance while random forest and XGBoost only have important features.

Table S1. **Numeric machine learning classification summary on both graphene-assisted and no-graphene Raman spectra.**

| Model | Graphene | Area | Accuracy | AUC | Sensitivity | Specificity |
|---|---|---|---|---|---|---|
| CatBoost | Yes | Cortex | 0.94 | 0.94 | 0.93 | 0.95 |
| CatBoost | Yes | Hippocampus | 0.68 | 0.69 | 0.70 | 0.68 |
| CatBoost | Yes | Thalamus | 0.80 | 0.80 | 0.75 | 0.85 |
| Random Forest | Yes | Cortex | 0.96 | 0.96 | 0.95 | 0.98 |
| Random Forest | Yes | Hippocampus | 0.72 | 0.72 | 0.70 | 0.74 |
| Random Forest | Yes | Thalamus | 0.88 | 0.88 | 0.87 | 0.89 |
| SVM | Yes | Cortex | 0.98 | 0.98 | 0.99 | 0.98 |
| SVM | Yes | Hippocampus | 0.71 | 0.70 | 0.75 | 0.65 |
| SVM | Yes | Thalamus | 0.96 | 0.96 | 0.93 | 0.98 |
| XGBoost | Yes | Cortex | 0.93 | 0.93 | 0.93 | 0.93 |
| XGBoost | Yes | Hippocampus | 0.68 | 0.68 | 0.70 | 0.65 |
| XGBoost | Yes | Thalamus | 0.83 | 0.83 | 0.81 | 0.86 |
| CatBoost | No | Cortex | 0.77 | 0.77 | 0.76 | 0.78 |
| CatBoost | No | Hippocampus | 0.81 | 0.81 | 0.80 | 0.82 |
| CatBoost | No | Thalamus | 0.75 | 0.75 | 0.75 | 0.76 |
| Random Forest | No | Cortex | 0.77 | 0.79 | 0.76 | 0.80 |
| Random Forest | No | Hippocampus | 0.92 | 0.92 | 0.90 | 0.94 |
| Random Forest | No | Thalamus | 0.79 | 0.78 | 0.77 | 0.79 |
| SVM | No | Cortex | 0.70 | 0.70 | 0.72 | 0.68 |
| SVM | No | Hippocampus | 0.86 | 0.87 | 0.86 | 0.88 |
| SVM | No | Thalamus | 0.87 | 0.88 | 0.87 | 0.89 |
| XGBoost | No | Cortex | 0.65 | 0.66 | 0.64 | 0.68 |
| XGBoost | No | Hippocampus | 0.78 | 0.79 | 0.80 | 0.78 |
| XGBoost | No | Thalamus | 0.77 | 0.77 | 0.78 | 0.76 |

Table S2. **17 major composite molecules of the brain and their corresponding literatures.**

| Component Molecule | Reference |
|---|---|
| Actin | [1] |
| Alpha-synuclein | [2,3] |
| Aspartate Aminotransferase | [4] |
| Beta-synuclein | [3] |
| Cholesterol | [5] |
| Cytochrome | [6] |
| DNA | [1] |
| Glycogen | [7] |
| Hemoglobin Alpha | [8] |
| Hemoglobin Beta | [8] |
| Lactate Dehydrogenase | [9] |
| Myelin Basic Protein | [10] |
| Phosphatidylcholine | [11] |
| RNA | [12] |
| Triolein | [1,13] |
| Tropomyosin | [14,15] |
| Ubiquitin | [16] |
